# Spatiotemporal analysis for detection of pre-symptomatic shape changes in neurodegenerative diseases: applied to GENFI study

**DOI:** 10.1101/385427

**Authors:** Claire Cury, Stanley Durrleman, David Cash, Marco Lorenzi, Jennifer M Nicholas, Martina Bocchetta, John C. van Swieten, Barbara Borroni, Daniela Galimberti, Mario Masellis, Maria Carmela Tartaglia, James B Rowe, Caroline Graff, Fabrizio Tagliavini, Giovanni B. Frisoni, Robert Laforce, Elizabeth Finger, Alexandre de Mendonca, Sandro Sorbi, Sebastien Ourselin, Jonathan D. Rohrer, Marc Modat, on behalf of the Genetic FTD Initiative, GENFI

## Abstract

Brain atrophy as measured from structural MR images, is one of the primary imaging biomarkers used to track neurodegenerative disease progression. In diseases such as frontotemporal dementia or Alzheimer’s disease, atrophy can be observed in key brain structures years before any clinical symptoms are present. Atrophy is most commonly captured as volume change of key structures and the shape changes of these structures are typically not analysed despite being potentially more sensitive than summary volume statistics over the entire structure.

In this paper we propose a spatiotemporal analysis pipeline based Large Diffeomorphic Deformation Metric Mapping (LDDMM) to detect shape changes from volumetric MRI scans. We applied our framework to a cohort of individuals with genetic variants of frontotemporal dementia and healthy controls from the Genetic FTD Initiative (GENFI) study. Our method, take full advantage of the LDDMM framework, and relies on the creation of a population specific average spatiotemporal trajectory of a relevant brain structure of interest, the thalamus in our case. The residuals from each patient data to the average spatiotemporal trajectory are then clustered and studied to assess when presymptomatic mutation carriers differ from healthy control subjects.

We found statistical differences in shape in the anterior region of the thalamus at least five years before the mutation carrier subjects develop any clinical symptoms. This region of the thalamus has been shown to be predominantly connected to the frontal lobe, consistent with the pattern of cortical atrophy seen in the disease.

## 1. Introduction

Neurodegenerative diseases such as frontotemporal dementia (FTD) present with progressive symptoms of behavioural and cognitive dysfunction. These changes follow many years of a clinically silent phase in the disease, where abnormal protein pathology slowly accumulates within the brain, leading to neurodegenerative processes that ultimately result in loss of function. Reliably identifying presymptomatic changes in individuals could lead to intervention with therapies that could slow, or even halt, the onset of these diseases. However, finding a cohort of presymptomatic individuals guaranteed to develop a form of dementia can be challenging. One common strategy is to investigate people who are at-risk for rare autosomal dominant forms of dementia. Half of these individuals are carriers of the mutation, allowing for comparisons between carriers and non-carriers at various stages within the disease process. In the case of genetic FTD, roughly one third of all cases are caused by autosomal dominant mutations, primarily in three genes: chromosome 9 open reading frame 72 (*C9orf72*), progranulin (*GRN*), and microtubule associated protein tau (*MAPT*) [1]. As the name would suggest, in all mutations, there is early involvement of both the frontal and temporal lobes, as well as the insula where differences can be observed as early as ten years before estimated age of expected symptom onset, as shown in Rohrer *et al*. [2]. However, there are additional structures, such as the thalamus, which also appear to be implicated to some degree early on in the disease process [3]. In many forms of FTD, clinical presentations suggest a left/right asymmetry in terms of which hemisphere is more affected, and this is often supported by evidence of increased atrophy within the affected hemisphere [4]. However, the affected side is not consistent across all cases, and in some cases, there is no evidence of an asymmetry. As this asymmetry is likely to start early in the disease process, it must be taken into account when looking to detect early changes with any sensitivity.

One biomarker that shows promise during the presymptomatic phase is measurement of atrophy derived from structural magnetic resonance imaging (**MRI**) [5, 2, 6] Volumes summarizing change within a region of interest (**ROI**) tend to be more sensitive to early change than voxelwise approaches, but they do not provide any spatial localisation as to where the atrophy is occurring within the **ROI**. Conversely, voxelwise analysis can provide better spatial localisation, but the mass univariate nature of the analysis requires correction for multiple comparisons to control for false positive findings, which often results in reduced sensitivity. As loss of brain volume will imply a change in the shape of the structure, a third option is to perform the shape analysis over time for a structure of interest. This could provide more spatial information than a single summary measure of volume alone, but does not require the same level of multiple comparisons as a voxelwise analyses. Given the decades long nature of the disease process, it is not yet feasible to measure the complete time course within one individual. Therefore, the pattern of atrophy over the course of the disease must be estimated through spatiotemporal regression models based on large populations of either cross-sectional data or through longitudinal data that covers a smaller segment (i.e. a few years) of the disease process within each individual.

There have been numerous approaches to spatiotemporally model trajectories of ageing and dementia. Some methods model this evolution using dense 4D deformation fields to measure change between timepoints. Lorenzi *et al*. [7] modelled the 4D deformation fields within a population to obtain subject-specific measurements of atrophy. An extension of this work discriminated spatiotemporal patterns that could be attributed to natural ageing versus to those that were related to disease [8]. Other groups establish point correspondences between subjects on a surface representation, and then apply mixed effects models at those points [9, 10, 11], providing fixed effects that represent the change across the overall population while allowing individual longitudinal trajectories as random effects. Using more complex representations of surfaces, Durrleman et al. [12] proposed a spatiotemporal regression approach to estimate continuous subject-specific trajectories of longitudinal data.

In our previous work [13], we defined the shape of the structure of interest as its 3D outline that is rotation and translation invariant. Differences between shapes were quantified using the Large Deformation Diffeomorphic Metric Mapping (**LDDMM**) framework [14, 15, 16], producing a smooth and invertible continuum between all possible shapes within the population. The smooth representation of these deformations also acted as low-pass filter, reducing the effects of irregularities and errors in the surface boundaries. Overall, our approach consisted of three main steps. First, using all available data, we compute an average shape spatiotemporal trajectory. Second, for every individual shape we evaluate its distance from the mean trajectory. Last, after spatially normalising all the subject-specific distances to the mean, we run a statistical analysis on the subject-specific residuals to assess when a shape starts diverging from normality. This previous work presented a global spatio-temporal analysis, on one side of the brain, without considering a potential asymmetry of the disease. In this paper, we build on the aforementioned framework, which we altered in two main ways. First, we take into consideration the potential asymmetry of **FTD** by considering the left and right structures using a common shape representation. Second, we modified our feature extraction method using a clustering approach to ensure we can attribute the recovered differences to substructure of the shape under study, and made a novel local analysis, based on cluster of deformations, taking better advantage of the **LDDMM** framework.

We apply this approach to data from the Genetic **FTD** Initiative (**GENFI**), an international study of autosomal dominant forms of **FTD** aimed at collecting multimodal neuroimaging, alongside other biomarkers with the objective of obtaining an improved understanding of the changes that are occurring during the presymptomatic phase of the disease. In general, the expected age of onset of clinical symptoms is estimated by using the average age of onset in the family of the subject, allowing to align the different subjects onto a single time axis. We applied our method to a subcortical structure, the thalamus, which has been shown to present volumetric differences before onset in Rohrer *et al*. [2]. We used the expected age to onset to characterise the time progression. In the next section, we will present the different steps of the proposed framework before then further describing the experiment and associated results.

## 2. Method

We indicate with {(*S_i_*, *t_i_*)}_i∈{0;…;*N*–1_ a set of *N* shapes associated with a corresponding time point *t_i_*. With analogy to classical random-effect-modelling approaches, we assume that each shape is a random realisation of a common underlying spatiotemporal process *φ*(*t*):

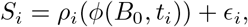

where *B*_0_ is a common reference frame, and *ρ_i_* is a subject-specific ”residual” deformation accounting for individual deviation from the mean shape. We characterise this residual through the diffeomorphism linking the shape *S_i_* to the corresponding sample of the common spatiotemporal trajectory at time point *t_i_*. We also assume that *ϵ_i_* is Gaussian randomly distributed noise. In order to identify group-wise differences between the given populations, we rely on the analysis of the subjects-specific residuals deformations *ρ_i_*.

This is a challenging problem, since all *ρ_i_* are defined at different time points along the common spatiotemporal trajectory, and therefore cannot be directly compared in a common anatomical framework. Moreover, the optimisation of the functional for the simultaneous estimation of the group-wise trajectory and random effects is not trivial, and would ultimately result in expensive and thus impractical numerical schemes. For these reasons, we propose a serial optimisation of the problem by introducing an efficient numerical framework composed of three steps illustrated in Figure 1.

(i) First, we assume that the residuals deformations *ρ_i_* are fixed, and we estimate the common trajectory *ϕ*(*t*). (ii) Second, given the modelled trajectory *ϕ*, we estimate the residuals deformations *ρ_i_* through non-linear registration between the trajectory point *ϕ*(*B*_0_,*t_i_*) and *S_i_*. (iii) Third, we spatially normalise the residual deformations in the common initial reference space *B*_0_ using parallel transport.

The proposed framework relies on the mathematical setting of the Large Diffeomorphic Deformation Metric Mapping (LDDMM) framework and the varifold representation of shapes (section 2.1). This choice allows a mathematically consistent definition of all steps (section 2.2), namely: (i) the spatiotemporal regression, (ii) the *ρ_i_* deformations estimation, and (iii) the normalisation of the initial momentum of *ρ_i_* through parallel transport.

**Figure 1:**
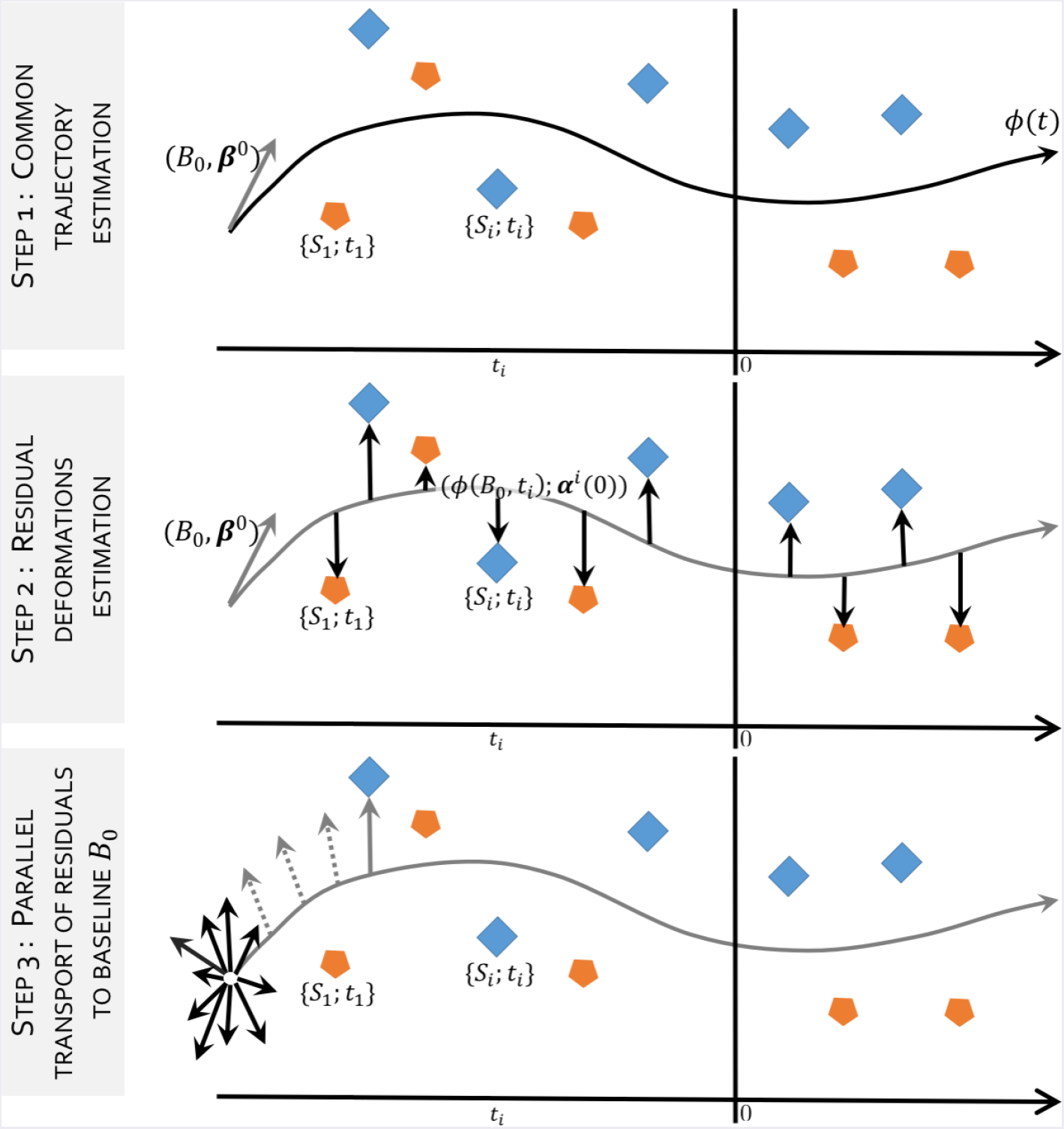
Overview of the proposed regression approach. The temporal axis indicates the time variable attached to the data. The residual deformations (step 2) *ρ_i_* parametrised by (*ϕ*(*B*_0_, *t_i_*); *α^i^*(0)) computed from the common trajectory (step 1) *ϕ* parametrised by (*B*_0_; *β*^0^), can not be analysed because they are defined on different spaces i.e. *ϕ*(*B*_0_,*t_i_*). They have to be transported to a common space (i.e. *B*_0_) along the geodesic *ϕ*, so they can be analysed (step 3).

### 2.1. Large diffeomorphic deformation metric mapping and varifold representation

The **LDDMM** framework [14, 15] is a mathematical and algorithmic framework based on flows of diffeomorphisms, which allows comparing anatomical shapes as well as performing statistics. The framework used in this paper is a discrete parametrisation of the **LDDMM** framework, as proposed by Durrleman *et al*. [17], based on a finite set of *N*_*B*_0__ control points overlaid on the 3D space enclosing the initial shape *B*_0_. The control points number and position are independent from the shapes being deformed as they do not require to be aligned with the shapes’ vertices. They are used to define a potentially infinite-dimensional basis for the parametrization of the deformation. Momentum vectors are associated with the control points and are used as weights for the decomposition of a given deformations onto this basis.

Deformation maps 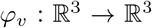 are built by integrating time-varying vector fields (*υ_t_*)_0≤*t*≤1_, such that each *υ*(·, *t*) belongs to a Reproducing Kernel Hilbert Space (**RKHS**) *V* with kernel *K_v_*. We use a Gaussian kernel for all control points *x, y*:

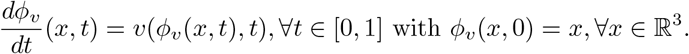

with Id the identity matrix, and λ a scale factor which determines the size of the kernel and therefore the degree of smoothness of the deformations. We define *φ*_*υ*_(*x*) = *ϕ*_*υ*_(*x*, 1) as the diffeomorphism induced by *υ*(*x,t*) where *ϕ_ν_* (*x*, 1) is the unique solution of the differential equation:

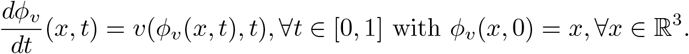

Velocity fields *υ*(·, *t*) are controlled via an energy functional 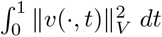 is a Hilbert norm defined on vector fields of 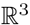, which is used as a regularity term in the matching functional to penalises non-regularity. In the **LDDMM** framework, matching two shapes *S* and *T* requires estimating an optimal deformation map 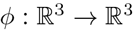 such that *ϕ*(*S*) is close to *T*. This is achieved by optimising

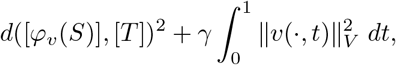

where γ balances the regularity of *ϕ_υ_* against the spatial proximity *d*, a similarity measure between the varifold representation of *φ_υ_*(*S*) and *T* noted respectively [*φ_υ_*(*S*)] and [*T*].

In a discrete setting, the vector fields *υ*(*x, t*) corresponding to optimal maps are expressed as combinations of spline parametrised fields that involve the reproducing kernel *K_v_* of the space *V*:

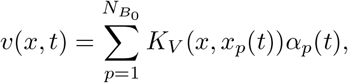

where *x_p_*(*t*) = *ϕ_υ_*(*x_p_, t*) are the trajectories of control points *x_p_*. The control points are regularly spaced on a 3D grid overlaid on the space that contains the mesh of the subject *S*. The control point spacing is defined by the size of the kernel *K_v_*. The time-dependent vectors 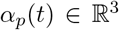 are referred to as momentum vectors attached to *x_p_*. The full deformation can be encoded by the set of initial momentum vectors *α*(0) = {*α_p_*(0)}_1≤*p*≤*n*_ located at the points {*x_p_*}_1≤*p*≤*n*_. This allows to analyse the set of deformation maps from a given template to the observed shapes by performing statistics on the initial momentum vectors defined on control points located around the template shape. The process of generating back any deformation maps from initial conditions (*x_p_*(0), *α_p_*(0)), i.e. integrating the geodesic equations, is called geodesic shooting or exponential map and is noted exp_*x_p_*(0)_(*α_p_*(0)).

As previously stated, varifolds are used to represent shapes [18]. They are nonoriented versions of the representation with currents [19], which are used to efficiently model a large range of shapes. To represent a shape *S* as a varifold, the shape space is embedded into the dual space of a Reproducing Kernel Hilbert Space (**RKHS**) W, noted *W*\*, and encoded using a set of non-oriented unit normals attached on each vertices of the shape. This kernel-based embedding allows to define a distance between different embedded shapes. Varifolds are robust to varying topologies, do not require point to point correspondences, and embed the shapes in a vector space, which facilitate the interpretation of results. The varifold representation of a discretised mesh composed by *M* triangles *S* is noted [*S*] and writes: 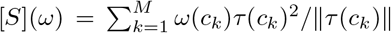 with *ω* a vector field in W, *c_k_* the centre of the triangle *k*, and *τ*(*c_k_*) the tangent of the surface *S* at point *c_k_*.

### 2.2. Residual extraction framework

Due to the asymmetry of the disease, the proposed framework has been designed so that it is unbiased to the affected side. For each subject, rather than considering the left or right structure, we build a mean shape by averaging both sides. First, we flip all input T1w brain images, in order to have all structures, left and right, on the same side, right. Second, we affinely align the T1w brain images (the original and the flipped once) to a subject-specific mid-space [20] before rigidly refining the alignment of the structure of interest, that has been segmented using the method proposed by Cardoso *et al*. [21]. Third, we extract the meshes of the left (flipped, *L_i_*) and right structures (*R_i_*), and compute the mean shape, by estimating the diffeomorphisms 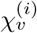 for each subject *i*, such as 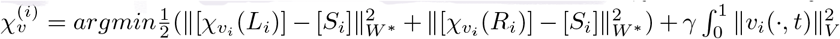 with *S_i_* the mean shape of subject *i* and *W*\* the space of varifolds. The obtained subject-specific average shape of the structure of interest is noted *S_i_* and is associated with a temporal information *t_i_*, the number of years to the expected onset (**EYO**) of the subject *i*.

The computation of the spatiotemporal regression [12] requires an initial shape *B*_0_ = {*x_P_*}_*P*=1,…,*N*_*B*_0___ as reference. To avoid any bias towards a subject selected as the initial shape, we estimate the initial shape from the 10 subjects who are the furthest away from expected symptom onset, so located in time around -40 years before EYO. We estimate the centroid of those 10 subjects using the diffeomorphic Iterative Centroid method [22], which estimate a centre of a given population in a reasonable computation time [23].

The spatiotemporal regression of the set of shapes {(*S_i_, t_i_*)}_*i*∈{0;…;*N*−1}_ is implemented in the Deformetrica software [24, 25]^2^. The EYO values are discretised into *T* time points. Starting from *B*_0_ at time *t* = 0, a geodesic moving through the positions *ϕ*(*B*_0_, *t*), ∀*t* ∈ {0;…; *T*} is computed by minimising the discrepancy between the model at time *t* (i.e. *ϕ*(*B*_0_, *t*)) and the observed shapes *S_i_*:

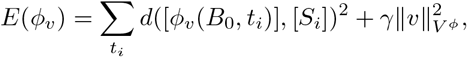

with *υ* the time-varying velocity vector field that belongs to the RKHS *V* determined by the Gaussian Kernel *K*. The initial momentum vectors 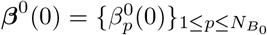 is defined on the control point grid overlay on the baseline shape *B*_0_ and fully encodes the geodesic regression parametrised by {*B*_0_; *β*^0^(0)}.

We then compute the residuals diffeomorphic deformations *ρ_i_* between every obser-vation and the spatio-temporal average shape by estimating a geodesic between *ϕ*(*B*_0_,*t_i_*) and {*S_i_, t_i_*}. This yields a set of trajectories parametrised by {*ϕ*(*B*_0_, *t_i_*); ***α**^i^* (0)}_*i*∈{0;…;*N*−1_} that encodes the deformations ρ_i_ from the spatio-temporal regression to all subjects, with ***α**^i^*(0) the initial momentum vectors, where the varying parameter is the step of the deformation. This should not be confused with the time we used until now which corresponds to EYO and time varying deformation of the main spatio-temporal trajectory.

In order to be able to compare this set of momenta, we gather them in the same Euclidean space. This is achieved by transporting all momenta into the initial space of *B*_0_ = *ϕ*(*B*_0_, 0), using a parallel transport method based on Jacobi fields as introduced in [26]. Parallel transporting a vector along a curve (the computed trajectory parametrised by (*B*_0_; *β*^0^(0))) consists in translating it across the tangent spaces along the curve by preserving its parallelism, according to a given connection. The Levi-Civita connection is used in the **LDDMM** framework. The vector is parallel transported along the curve if the connection is null for all steps along the curve [27]. We use Jacobi field instead of the Schild’s Ladder method [28], to avoid the cumulative errors and the excessive computation time due to the computation of Riemannian Logarithms in the **LDDMM** framework, required for the Schild’s Ladder. The cumulative errors would have differed from subject to subject and thus introduce a bias. Indeed, their distances from the baseline shape vary, as they all are at different points along the temporal axis. The Jacobi field, used to transport a vector ***α**^i^*(0) from a time *t* to the time *t*_0_ = 0 along the geodesic *γ*, is defined as:

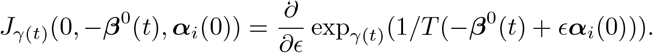

The transported initial momentum vector ***α**_i_*(0) is noted ***θ**_i_*(0). After parallel transporting all residuals, all initial momentum vectors are defined in *B*_0_.

### 2.3. Feature extraction for statistical analysis

Each transported initial momentum vectors ***θ**_i_*(0) is of size 3 × *N*_*B*_0__, where *N*_*B*_0__ is the number of control point used to parametrise the geodesics.

Jacobian determinants are commonly used to study shrinkage or growth of the surface, and are a geometric measure derived from the full deformation tensor. In this work we propose an analysis framework where we decouple the amplitude and the orientation of the deformation. Such approach still analyse growth and shrinkage, but also other geometric aspects, such as rotation and torsion, not captured by the surface Jacobian.

To analyse direct measures from deformation and to avoid losing statistical power from doing a large number of comparisons, we propose an original clustering by grouping the parametrisation (*B*_0_; *β*^0^(0)) of the spatio-temporal regression *ϕ* into clusters.

To do so, we defined a similarity measure derived from the positions of the control points *x_p_*, the pairwise angles and the magnitudes of the initial momentum vectors 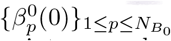 attached to the control point *x_p_*. The difference between two control points *x_p_* and *x_q_* ∀*p, q* ∈ {1;…; *N*_*B*_0__} is defined by the euclidean distance, the angle between two vectors is defined by the cosine. The similarity between *p* and *q* is defined by 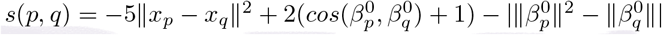. Parameters are chosen to balance between vector similarity and control point positions and depend on the distance in mm between two points. The distance is determined by the kernel *K_v_* so that clusters encompass control points and their momentum vectors within the same area and look alike. To estimate those clusters, we used a spectral clustering method [29] using the discretisation approach presented in [30] for initialisation, as it has been shown to be more stable than other approaches such as k-means for initialisation. 3000 different initialisations are generated and we select the best one in term of inertia for spectral clustering. We chose 10 clusters as thought this would be a good balance between reducing the number of multiple comparisons while maintaining some spatial specificity in the analyses and equitable clusters. A mean vector is then computed from the parallel transported residuals defined on the control points of the cluster. This is done for each cluster and for each subject. We then obtain N vectors {*ν_i,k_*} per cluster *k*, and 10 vectors per subject *i*.

For the statistical analysis, we will use two uncorrelated descriptors for the vectors {*ν_i,k_*}: the amplitude and the orientation. The orientation of the vectors {*ν_i,k_*} is originally represented by 3 angles, one per axis. The angles are then projected via a Principal Component Analysis on the first eigenvector, therefore the orientation of {*ν_i,k_*} consid-ered here is represented by one continuous scalar, leading to the set of responsive variable {*O_i,k_*}.

## 3. Data and application

As previously mentioned, we applied the proposed framework to the GENFI study and used the thalamus as structure of interest.

### Dataset description

All participants included in this study come from the data freeze 1 of the GENFI cohort described in detail in [2]. Initial results from this cohort [2] show volumetric differences in the thalamus at least 5 years before expected age of onset with an effect in all genetic subtypes, and so we chose this well-defined subcortical structure for further analysis. In this paper we used 211 participants, 113 mutation carriers (MAPT=26, GRN=53, C9ORF=34) and 98 non-carriers. All participants have a T1-weighted (T1w) MRI available and an associated expected years to symptom onset (EYO). The EYO is calculated as the difference between the age of the participant at the time of the T1w acquisition and the mean age at onset of affected family members, EYO range from -40 years to +20 years. Table 1 shows the demographics of the study participants used in this analysis.

### Application to the thalamus

T1w brain images of all subjects were affinely group-wise registered [20], before applying a rigid registration focused solely around the structure of interest. We then extracted the meshes corresponding to the thalamus, including around 2, 300 vertices. This resulted in 211 thalamus meshes, representing the mean left and the right shape. Each were associated with the EYO of the corresponding subject as well as mutation status: non-carrier and mutation carrier (MC). For the spatiotemporal regression, we used 30 time points, which corresponds approximatively to one time point every two years. The space of deformations V was defined using a 11mm width kernel, approximately half of the length of the thalamus, which leads to a set of 288 control points. For the space of varifolds we used a 5mm width kernel.

**Table 1:** Data demographics, in absolute values.

|  | Non-carriers<br>n=98 | Mutation carriers<br>n=113 |
| --- | --- | --- |
| Males | 59 | 56 |
| Asymptomatic | 98 | 76 |
| Age in years (med (IQR)) | 50.2 (36.6 - 62.1) | 52.7 (41.1 - 62.7) |
| Years from expected onset: |  |  |
| $\leq -20$ years | 30 | 21 |
| $-20 \leq \text{years} \leq -10$ | 16 | 21 |
| $-10 \leq \text{years} < 0$ | 23 | 22 |
| $0 \leq \text{years}$ | 29 | 49 |

Similarly to the volumetric analysis performed by Rohrer *et al*. [2], we used a mixed effect model to study the shape difference between the non-carriers and mutation carriers. Amplitude {|*ν_i k_*|} and orientation {*O_i k_*} were used as responsive variables and the fixed effects predictors of interest were mutation carrier status, EYO, interaction between mutation carrier status and EYO, sex and the site in which the subject has been scanned. A random intercept for family allows values of the marker to be correlated between family members.

We performed a Wald test for every model, assessing the difference between the mutation carrier group and the non-carrier group, and the evolution of differences across time. For each analysis with statistically significant differences between both groups, further Wald tests were conducted every 5 years as in the volumetric analysis [2] to assess how long before the expected onset we could detect changes between mutation carriers and controls.

## 4. Results

Results for the amplitude and the orientation of the residual momentum vectors are presented Table 2. We found significant differences, after correction for multiple comparisons, in cluster 1 and cluster 4, for both tests; T1:differences between MC and controls and T2: differences over time between MC and controls. Those differences are significant after Bonferroni correction for multiple comparisons (20 tests). Cluster 1 shows differences in the orientation, and no differences in the amplitude, whereas cluster 4 shows significant differences for those 2 tests in amplitude, and no differences in orientation. Those 2 clusters are thus selected for the next wald test step. Wald tests were conducted every 5 years between 20 years before the expected onset and 10 years after the expected onset to limit the number of tests, since we don’t expect changes before -20 EYO, and results are shown in Figure 2, the p-values and confidence intervals are corrected for multiple comparison across time using Bonferroni correction.

**Table 2:** p-values with the corresponding *χ*^2^ value, resulting from the Wald tests testing the mutation carrier (MC) differences (test T1), and the evolution of those differences along time (test T2), for the amplitude of the initial momentum vector and its orientation, for the clusters showing at least one significant test. Bold p-values: ≤ 0.05, and starred (*) p-values indicate the corrected threshold for multiple comparisons: ≤ 2.5e-3.

|  |  | C1 | C2 | C4 | C6 | C7 |
| --- | --- | --- | --- | --- | --- | --- |
| Ampl. | T1 | $p = 0.48$ | $p = 0.51$ | $p = \mathbf{1.5e-3 (*)}$ | $p = 0.08$ | $p = 0.76$ |
| | | $\chi^2_{df=2} = 1.43$ | $\chi^2_{df=2} = 1.35$ | $\chi^2_{df=2} = 12.94$ | $\chi^2_{df=2} = 5.10$ | $\chi^2_{df=2} = 0.55$ |
| | T2 | $p = 0.24$ | $p = 0.26$ | $p = \mathbf{1.5e-3 (*)}$ | $p = \mathbf{0.04}$ | $p = 0.68$ |
| | | $\chi^2_{df=1} = 1.37$ | $\chi^2_{df=1} = 1.28$ | $\chi^2_{df=1} = 10.08$ | $\chi^2_{df=1} = 4.20$ | $\chi^2_{df=1} = 0.17$ |
| Orient. | T1 | $p = \mathbf{2e-4 (*)}$ | $p = 0.12$ | $p = 0.85$ | $p = 0.63$ | $p = 0.08$ |
| | | $\chi^2_{df=2} = 16.60$ | $\chi^2_{df=2} = 4.17$ | $\chi^2_{df=2} = 0.33$ | $\chi^2_{df=2} = 0.92$ | $\chi^2_{df=2} = 5.06$ |
| | T2 | $p = \mathbf{9e-4 (*)}$ | $p = \mathbf{0.05}$ | $p = 0.62$ | $p = 0.34$ | $p = \mathbf{0.04}$ |
| | | $\chi^2_{df=1} = 11.01$ | $\chi^2_{df=1} = 3.85$ | $\chi^2_{df=1} = 0.25$ | $\chi^2_{df=1} = 0.91$ | $\chi^2_{df=1} = 4.29$ |

The orientation of the cluster 1 deformation shows significant differences between the mutation carriers and controls, 5 years before EYO (*p* = 0.03), the uncorrected for this cluster is *p* = 2e -3, to keep a head to head comparison with the previous studies on this dataset [2, 13] in which the p-values at -5 EYO was significant but higher than here. The uncorrected p-values show significant differences at 10 years before EYO, with p=0.048 for the orientation of cluster 1. The amplitude between the two groups doesn’t differ significantly for the cluster 4 before EYO for corrected p-values, and differs 5 years before onset without correction (p=0.05). Figure 3 shows the initial momentum vectors of clusters 1 and 4, and the amount of displacement due to the deformations corresponding to those clusters 1 and 4, where each cluster has its own colour scale, since the maximum displacement for cluster 4 is about 3 mm, against 9 mm for cluster 1. Deformations affect more the anterior part of the thalamus.

Since the number of clusters used (10), is an arbitrary choice, we tried to reproduce the results with different number of clusters. We performed the analysis for 2, 4, 6, 8, 10, 12, 14 and 16 clusters. For 6 clusters and 16 clusters, there were differences in orientation for one of the clusters which deformation corresponds to the one of cluster 1 (see Figure 3). From 8 clusters to 14 clusters, we found a cluster with strong differences 5 years before the expected onset (*p* < 0.01) in orientation whose deformation corresponds again to the one of the cluster 1 (*p* = 0.003). The change in orientation for the deformation recovered within cluster 1 (see Figure 3) appears to be stable for different clusterings of the deformation parametrisation of the global spatiotemporal trajectory. All results regarding the different number of clusters can be found in supplementary material (`doi. org/10.5281/zenodo.1324234`).

## 5. Discussion and conclusion

We applied a novel method of statistical shape analysis to a cohort of individuals with genetic **FTD** in order to localise any presymptomatic differences present in the shape of the thalamus. From the analysis, we conclude that differences are observed five years before expected symptom onset. While volumetric analysis [2] and our initial shape analysis [13] also found these changes, this method showed significance that survived correction for multiple comparisons. The change in shape is primarily attributable to differences in orientation of the deformation rather than changes in amplitude of the deformation, which would imply a simple scaling effect of the region. This result confirms our previous shape analysis in this cohort [13] that was performed at a global level through a kernel principal component analysis. The first mode of variation which detected significant shape differences around the same point with respect to EYO did not capture volume differences but only changes in the orientation of the deformation. The results of those studies seem to indicate that shape changes occur before volume changes. As many regions of the thalamus contain a mixture of grey and white matter, these shape changes may reflect subtle shifts in the ratio between these two tissue types in the areas affected.

**Figure 2:**
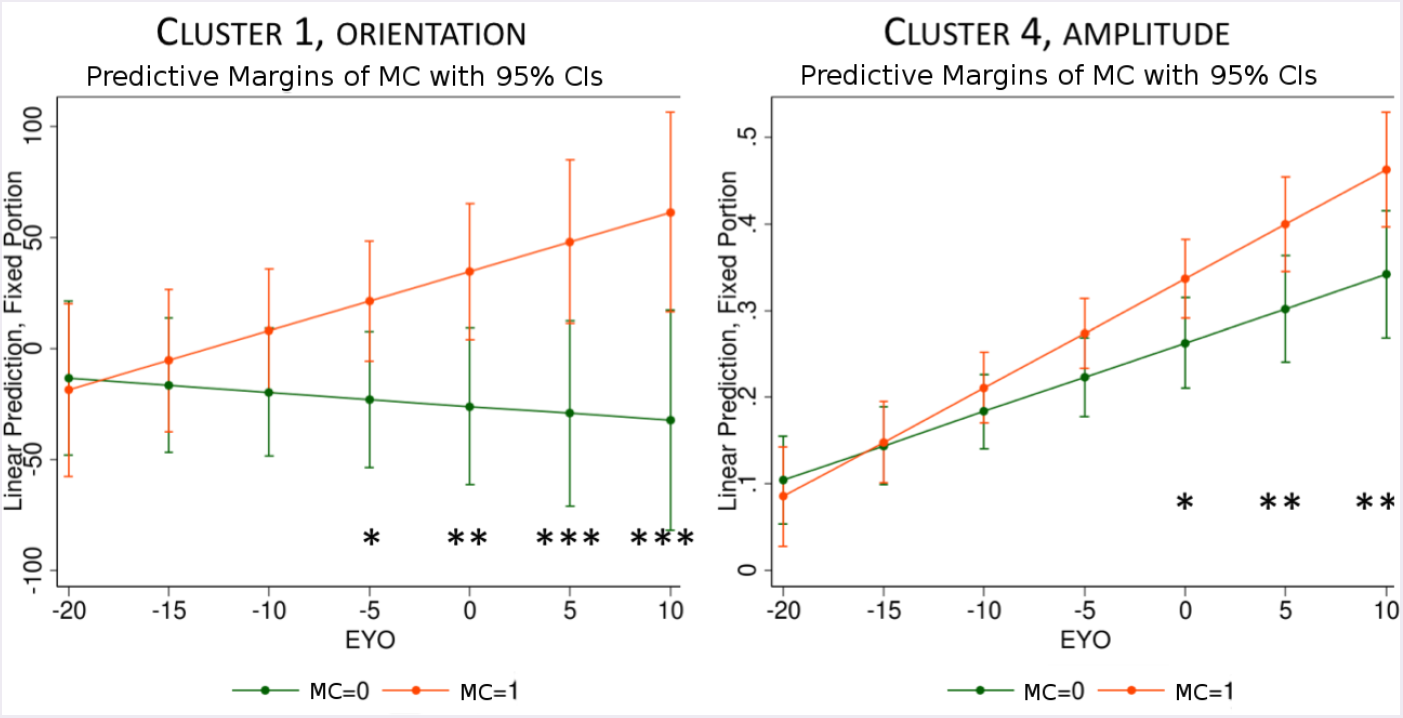
cluster 1 (orientation component) and cluster 4 (amplitude component) estimates in mutation carriers and controls, by estimated time from expected symptoms onset (EYO). p-values and confident interval are Bonferroni corrected. *: *p* < 0.05, **: *p* < 0.01, ***: *p* < 0.001

The regions of the thalamus most affected in the analysis are anterior, overlapping with the anterior nuclei group. The main connections of these nuclei are to the pre-frontal cortices, an area universally affected in all genetic forms of FTD. To illustrate this purpose, we used the Oxford thalamic connectivity atlas, a thalamic atlas based on its anatomical connectivity to the cerebral cortex [31], and displayed at Figure 4 the atlas next to the clusters 1 and 4. Whilst differences are seen in cortical involvement within the different genetic forms of **FTD** [32], it may well be that this joint analysis of GRN, C9orf72 and MAPT mutations is only identifying thalamic regions jointly affected.

Another interesting cortical region involved in FTD, could also be analysed with this method: the insula, which is located in the lateral sulci and is connected to the limbic system, and to the thalamus. It would be interesting to analyse the insula and thalamus together, and the insula only, so we could investigate if shape changes in both structures are linked.

**Figure 3:**
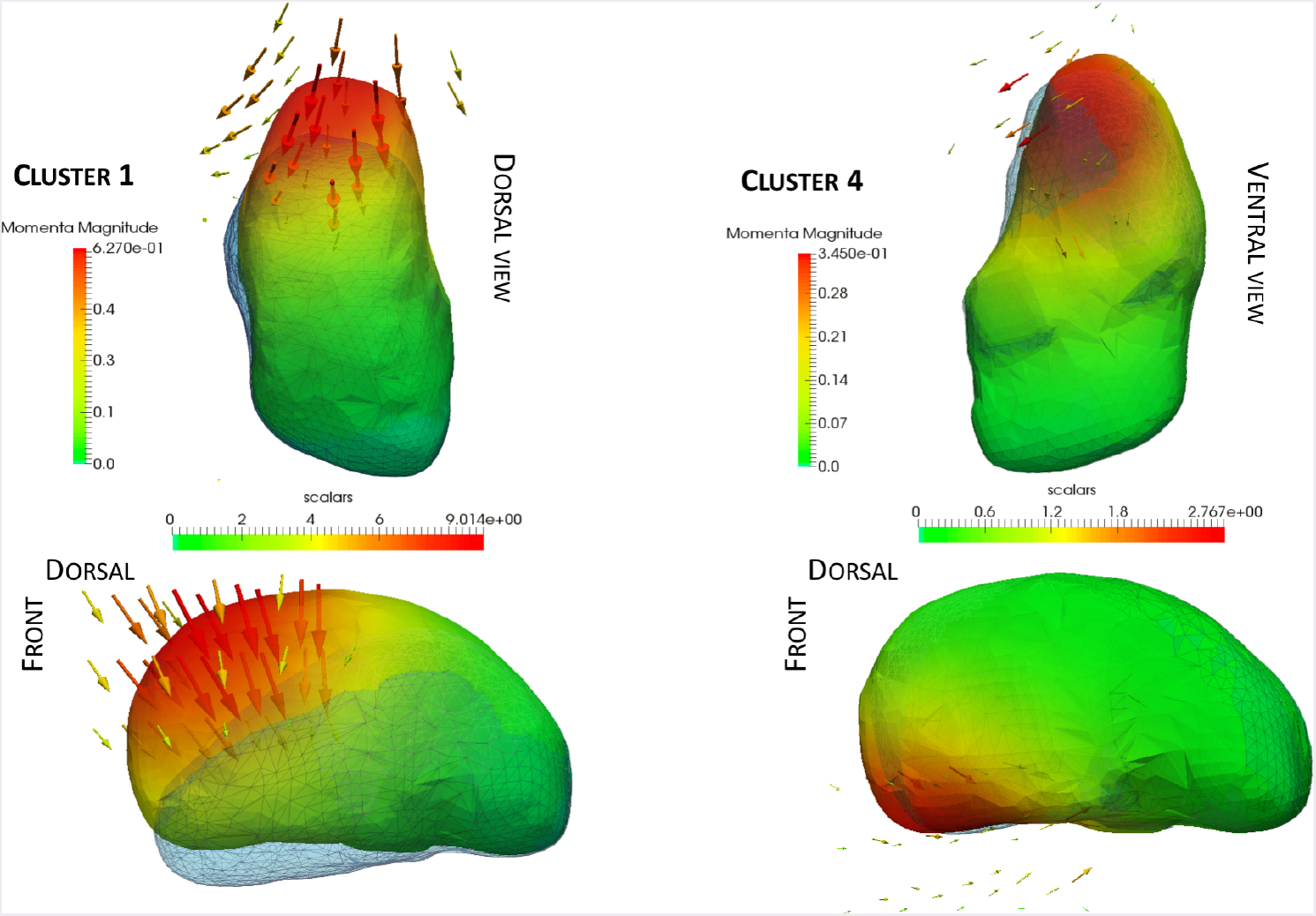
Deformation obtained by the momentum vectors (displayed here and coloured by amplitude) of Cluster 1 and Cluster 4. The colour map is in millimetres and indicates the displacement due to the corresponding deformation (blue meshes). The scale for Cluster 1 range from 0 mm to 9 mm, and from 0 mm to 2.8 mm for Cluster 4.

**Figure 4:**
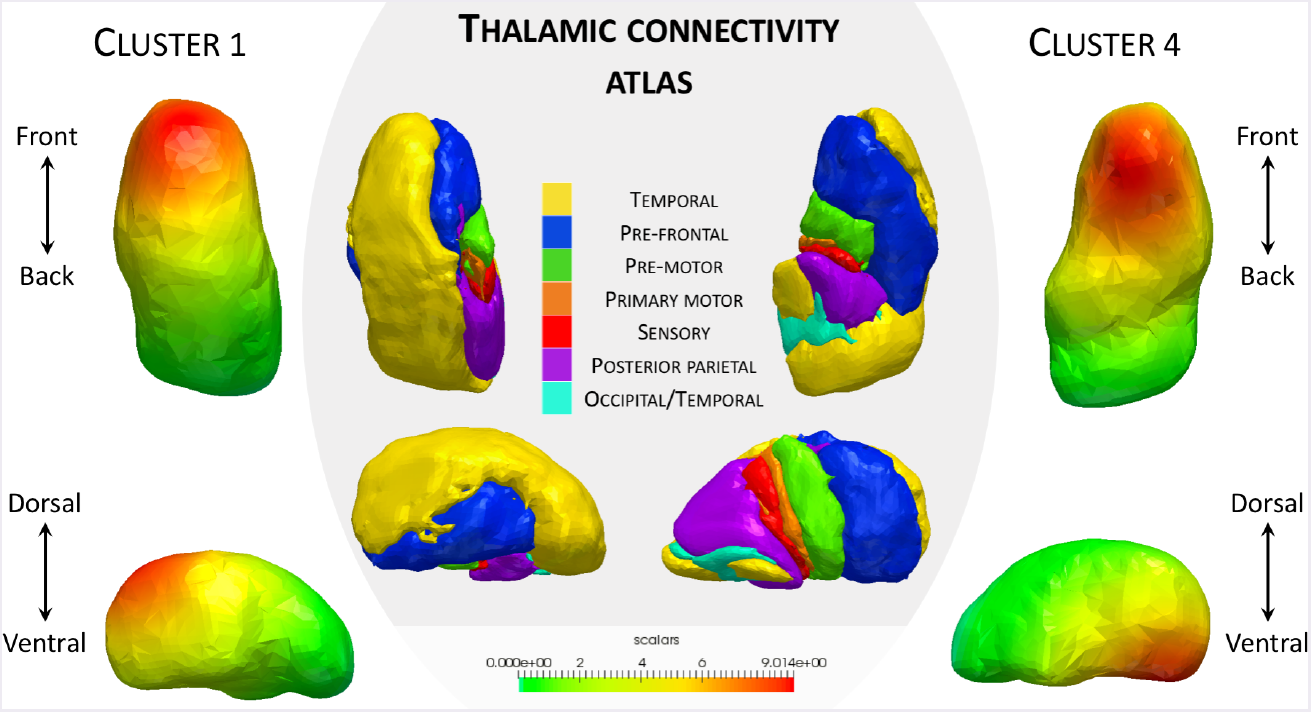
Thalamic connectivity atlas, and deformations clusters 1 and 4. The orientation of cluster 1 leads to significant differences between MC and controls 5 years before EYO.

The small numbers in each group precluded any analysis of the individual genetic types, but it will be important to investigate future data freezes from the GENFI study with larger numbers, particularly the C9orf72 group who have been shown to have early thalamic involvement [32].

Future studies should also evaluate the initial momentum vectors of individual geodesic evolution of shapes from each subject, through longitudinal data. Those individual evolutions will provide information on the differences of evolutions of shape between the mutation carriers and the controls.

## Acknowledgements

Claire Cury is supported by the EU-FP7 project VPH-DARE@IT (FP7-ICT-2011-9-601055). Stanley Durrleman has received funding from the program Investissements davenir ANR-10-IAIHU-06 and the European Unions Horizon 2020 research and innovation programme EuroPOND under grant agreement No 666992, and is funded by the European Research Council (ERC) under grant agreement No 678304. Marco Lorenzi received funding from the EPSRC (EP/J020990/1). Jennifer Nicholas is supported by UK Medical Research Council (grant MR/M023664/1). David Cash is supported by grants from the Alzheimer Society(AS-PG-15-025), Alzheimers Research UK (ARUK-PG2014-1946) and Medical Research Council UK (MR/M023664/1). JBR is supported by the Wellcome Trust (103838). Jonathan D. Rohrer is an MRC Clinician Scientist and has received funding from the NIHR Rare Diseases Translational Research Collaboration. Sebastien Ourselin receives funding from the EPSRC (EP/H046410/1, EP/K005278), the MRC (MR/J01107X/1), the NIHR Biomedical Research Unit (Dementia) at UCL and the National Institute for Health Research University College London Hospitals Biomedical Research Centre (NIHR BRC UCLH/UCL High Impact Initiative-BW.mn.BRC10269). Marc Modat is supported by the UCL Leonard Wolfson Experimental Neurology Centre (PR/ylr/18575) and Alzheimers Society UK (AS-PG-15-025). We would like to thank the participants and their families for taking part in the **GENFI** study.

## Footnotes

1 List of consortium members in appendix.

2 http://www.deformetrica.org/

